# CRISPR-Cas9 mediated knockout of SagD gene for overexpression of streptokinase in *Streptococcus equisimilis*

**DOI:** 10.1101/2021.10.20.465095

**Authors:** Armi M. Chaudhari, Sachin Vyas, Amrutlal Patel, Vijai Singh, Chaitanya G. Joshi, Madhvi Joshi

**Affiliations:** Gujarat Biotechnology Research Centre (GBRC), Department of Science and Technology, MS Building, 6th Floor, Sector- 11, Gandhinagar - 382011, Gujarat, India; Department of Biosciences, School of Science, Indrashil University, Rajpur, Mehsana 382715, Gujarat, India

**Keywords:** FasX, SagD, CRISPR-Cas9, Knockout, Streptokinase, therapy

## Abstract

Streptokinase is an enzyme that can break down the blood clots in some cases of myocardial infarction (Heart attack), pulmonary embolism, and arterial thromboembolism. Demand for streptokinase is high globally than the production due to increased incidences of various heart conditions. The main source of streptokinase is from various strains of *Streptococcus*. Expression of streptokinase in native strain *Streptococcus equisimilis* is limited due to the *SagD* inhibitor gene for production of streptokinase that needs to be knocked out in order to increase it expression. However, FasX is a small RNA (sRNA) present in group A *Streptococcus* species which is responsible for post-transcriptional regulation of streptokinase (*ska*) gene by binding at the 5’ end of *ska* mRNA. *S. equisimilis* is a *β*-hemolysin producing streptococcus bacterium (group C) containing the orthologue of FasX and natively expresses a clinically important thrombolytic streptokinase. In order to improve the stability of mRNA and increasing the expression of streptokinase which is inhibited by *SagD*. We used CRISPR-Cas9 to successfully knock-out of *SagD* gene and observed a 13.58-fold relative quantification of streptokinase expression in the mutant strain as compared to wild type. We have also demonstrated the successful target gene knockout using CRISPR-Cas9 in *S. equisimilis* that engineered strain can be used further for overexpression of streptokinase for therapeutic applications.

**Graphical Abstract:** 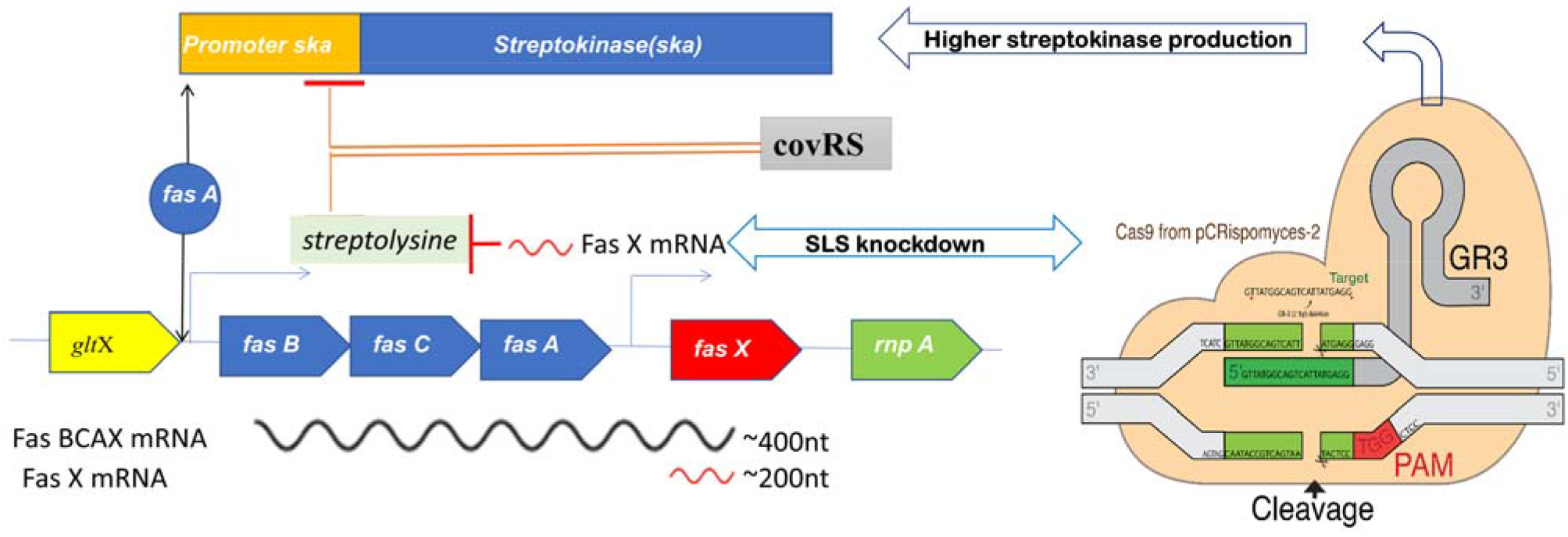

## 1. Introduction

Streptokinase, a ∼47 kDa thrombolytic enzyme that is naturally produced by a *β*-hemolytic group of *Streptococcus* species. Streptokinase forms a 1:1 streptokinase: plasminogen complex that is used as a blood clot buster in healthcare to save human life. Currently, increasing cases of myocardial infractions lead to the development of clot blusters with reduced hypersensitivity. Production of streptokinase in other host factors such as *Escherichia coli* has an issue of product toxicity or some other factors absence of small regulatory RNAs in expression host for the stability of streptokinase mRNA transcript [1,2]. Therefore, a pressing need has been arisen to engineer or genetic manipulate natural producer of streptokinase.

*Streptococcus* (Group A) possesses two-component regulatory systems [3] FasBCAX and CovRS [4]. Small regulatory ncRNAs (non-coding RNAs) from FasX stabilizes the mRNA transcript of *skc* that enhances the streptokinase production [2,5]. FasBCAX enhances the streptokinase production and also down-regulates the Streptolysin production, simultaneously CovRS up-regulates the Streptolysin production and down-regulates the streptokinase expression [2,4,6]. Therefore, Streptolysin produces by operon coding *SagABCDEFG* genes whereas, 2.7 kDa peptide (SagA) is undergone for post-translational modification by SagBCD complex which is similar to *E. coli* mcbBCD: B-Cyclohydratase, C-Dehydrogenase, and D-docking proteins and transported by SagEFG mediated ABC transporters [7,8]. A computer modeling approach was used to target the particular region of *SagD* gene for efficient downregulation of Streptolysin. The crystal structure was not determined yet for SagBCD complex, however, it has structural homology with *E*.*coli* MCB complex that used as template for generation of 3-dimensinal model [7,8]. It is useful for better understanding of its structure.

We hypothesized that the knockout of Streptolysin-producing genes which is under the regulation of CovRS may enhance mRNA stability for *skc* gene and also improve streptokinase production [4]. The direct target of SagA (which forms the Streptolysin peptide) knockout/knockdown was not performed as it is involved in cell survival and pathogenicity [9] rather another important factor was targeted to downregulate the Streptolysin. We have targeted the *SagD* gene to abrupt the Streptolysin post-translation modification, which may lead to enhance streptokinase production and decreases *Streptococcus* virulence [10,11]. In order to knock out of *SagD* gene, we used the CRISPR-Cas9 system which is widely used for targeted genome editing of several organisms [12–17]. In the present study, we aimed to increase the expression of streptokinase by CRISPR-Cas9 based knocking out of *SagD* gene in *S. equisimilis*.

## 2. Materials and Methods

### 2.1. Microorganisms, plasmids, and reagents

*S. equisimilis* culture used in this study was obtained from ATCC (**43079**.). *Escherichia coli* Top10F’ (Invitrogen, Thermo-Fischer Scientific, USA) was used for routine experimental work and also used for preparation of competent cells. The pCRISPomyces-2 vector used in this study was obtained from Addgene (Code: 617374). All the media and chemicals were used in this study were purchased from Hi-media (Mumbai-India) and Fischer Scientific (India). All the clones and cultures were stored in (25 % v/v) glycerol at -80 °C for long-term storage and further study.

### 2.2. Media and culture conditions

*S. equisimilis* was cultivated in Tryptonesoya broth (TSB) medium (HI media, Mumbai). *E. coli* Top10F’ strain was cultivated in Luria-Bertani (LB) medium (HI media, Mumbai) with an appropriate concentration of antibiotic (Apramycine) 50mg/L.

### 2.3. Design and construction of sgRNAs plasmids

The protospacer sequence [20 nucleotides (nt)], a complementary to target sequence *SagD* gene in *S. equisimilis* to construct sgRNA design was performed using the online available tool CRISPOR (http://crispor.tefor.net/) [18]. The genome of *S. equisimilis* was used by providing accession id to the tefor team to generate efficient gRNA sequences with minimum off-target effect. A total 1693 bp long target sequence for the *SagD* gene was used as an input. All of the 157 possible gRNAs sequences that can target PAM (Protospacer Adjacent Motif) sequences were analyzed based on efficiency and off-targets for mismatches. Four possible 24 nucleotides long protospacer sequences (4 nt 5’ sticky end + 20 nt spacer sequence) with the sticky ends ACGC on the forward sequence and AAAC on the reverse sequence were chemically synthesized [19]. The protospacer sequence was generated by equimolar ligation as previously reported (Supplementary Figure S1) [19]. The gRNA sequence containing sticky end overhands was inserted into CRISPomyces-2 plasmid using golden-gate assembly protocol using Bbs1 (type IIs restriction enzyme) (Supplementary Figure S1) [20].

### 2.4. Design of genome editing template for *SagD* gene knockout

Our major approach for design of genome editing template was a complete deletion of *SagD* gene region from *Streptococcus* genome. It was known that to overcome Cas9 mediated double-stranded endonuclease break, the prokaryotic system uses both repair mechanism either homologous end-joining pathway (NHEJ) or homology recombination(HR) pathways [21,22]. We synthesized the genome editing template using the overlap extension PCR method. For SagC, forward primer OEPSagCF 5’TATCTGAGAAGCTCGCAGAC 3’and reverse primer OEPSagCR 5’ <u>CCTTAGACCTCCTTATGACT</u>CTTCTTGAAGGAG 3’ were and SagE forward primer OEPSagEF 5’ <u>AGTCATAAGGAGGTCTAAGG</u>ATGCCTTTGTACACCCAATG 3’ and OEPSagER 5’ AGCACAGGCTGGTGATCAAC 3’. The size of PCR product for both genes SagC and sagE were 500 bp. OEFPSagCR and OEPSagEF shares overlapping complementary region which is underlined. Two PCR reactions were performed for SagC and SagE. PCR conditions were initial denaturation 95 °C 5min (1 cycle), 95 °C 30sec, 55 °C 15sec, 72 °C 1min (35 cycles), and last final extension stage 72 °C 5min (1 cycle) following 4°C. PCR products of both *SagC* and *SagE* were loaded into 1% agarose gel and we have eluted with manufacturer protocols. The Agarose elution step was to get rid of these four primers present along with PCR products. The presence four primers were hindered overlap extension PCR. Eluted products of *SagC* and *SagE* were added with an equimolar concentration of 500ng/µL. Protocol for overlap extension PCR was 95 °C: 1min 60 °C: 1 min 72 °C: 1 min 72 °C used followed by 72 °C for 5 min extension and kept at 4 °C. After this editing template was created using final PCR with OEPSagCF and OEPSagER primers and 1 kb product was generated. This was further eluted using agarose gel extraction kit (QIAGEN), and it was used for electroporation.

### 2.5. Transformation of constructed plasmid into *E. coli* Top 10F’ and *S. equimilus*

The irreversible reaction containing only intact plasmids with correct protospacer insert at the end was used for transformation into chemically competent *E. coli* TOP10F’ (∼10-100 ng) using a heat shock method [20,23]. It was spread on LB agar containing Apramycine (50 µg/mL) pre-coated with appropriate concentrations of IPTG (**1mM**) and X-Gal (40µg/ml) for blue-white colony screening [24]. White colonies observed were screened for plasmid isolation and confirmation by PCR using previously designed primers [20]. The protospacer insertion was also confirmed using Sanger sequencing.

Recombinant plasmids with sRNA were confirmed and transformed into electrocompetent *S. equisimilis* using electroporation methods. The electrocompetent cells preparation was performed according to the previously reported protocol [25]. Briefly, ∼0.4 optical density (O.D.) cells previously grown in glycine medium were washed four times with pre-chilled 0.5 M sucrose. The cells were suspended in 250 μL of a 10% (V/V) glycerol and 0.5M sucrose solution and divided into aliquots and stored at -80 °C for future electroporation experiments. Electroporation conditions were as followed 25 kV/cm, 200 Ω, and 25 μF [26]. Following the electroporation experiment, the cells were grown in 1 ml of TSB broth medium at 30 °C for 1 hour. The transformed cells were then plated on a TSB medium containing apramycin at a concentration of 50 µg/mL and incubated at 28 °C for 72 hours.

### 2.6. Screening of knocked-out clones using sequencing and hemolysis activity

Each of the colonies observed was tested for presence of recombinant plasmid using the vector-specific forward primer 5’ACGGCTGCCAGATAAGGCTT3’ and reverse primer 5’ TTCGCCACCTCTGACTTGAG3. The positive colonies were further grown in 5 mL of 50 µg/mL apramycin containing a TSB medium at 28 °C and 120 rpm. The positive colonies were screened for DNA sequencing of the PCR product that was obtained by *SagD* gene-specific primers for possible modifications and knockout (supplementary table2).

### 2.7. Real-time PCR for gene-expression profiles

Total RNA extraction was performed from the wild-type and mutant clones using RNeasy plus mini kit (QIAGEN). RNA was quantified by qubit 4. The cDNA synthesis was performed using the QunatiTect Reverse transcription kit (QIAGEN) [27]. The cDNA concentration was quantified for both wild-type and mutant samples as such that one reaction have 100ng cDNA. RT-PCR (Fast 7500 of Applied Biosystems) was performed using TB-Green™ Premix Ex Taq™ II. *SagD* gene expression was checked with forward primer SagD_RT_F 5’ GACAGCCTCTCATACAACAC 3’ reverse primer SagD_RT_R 5’AGCGGATTATCCTCTCCAAC 3’, *skc* gene expression has been checked using forward primer SK_RT_F 5’ ATACATCTTGACGGGTCAGG 3’ and reverse primer SK_RT_R 5’ AAGAGACCCTGCTGCCAT 3’. DNA gyrase was used as an endogenous control. In the reaction step, 1^st^ stage was holding stage at 52 °C for 2 min, initial denaturation at 95 °C for 2 min and in cycling stage (40 cycles) 1^st^ step was at 95 °C for 30 seconds and annealing temperature at 58 °C for 30 seconds followed by melting curve stage [28]. Then expression profiles of both *SagD* and *Sks* genes were analyzed using RT-PCR and a comparison was made.

### 2.8. MALDI-TOF/MS for confirmation of streptokinase expression on SDS-PAGE

SDS-PAGE was performed using crude extract supernatant from overnight grown cultures for both wild-type and clones using prescribe protocol in Sambrook [29]. For the confirmation of proteins, the prominent expected bands for Streptokinase (Ska) were eluted from the gel by cutting and subjected for in gel trypsin digestion [30]. Eluted peptides were used for MALDI-TOF/MS analysis in Auto flex speed MALDI/TOF/TOF spectrophotometer (Bruker Daltonics, Germany). Output data was submitted to MOSCOT server (http://www.matrixscience.com/server.html) for peptide mass finger printing analysis. In another set of experiments, the SDS-PAGE was performed using 1% (w/v) substrate containing PAGE gel. Zymography analysis was used to confirm the presence of an enzyme.

### 2.9. Streptokinase blood clot lysis and fibrin plate assay

Crude enzyme supernatant was obtained from the culture of both wild-type and clones (mutants). It was subjected to clot lysis assay [31]. Wild-type and mutant strains were grown in TSB media overnight and in the next day, equal O.D. was set to 0.05 at A_600nm_. In brief, 10 ml of fresh blood sample was collected from a voluntary healthy individual (As per Institutional Animal Ethical Guideline). Further 500 µL of blood was collected and allowed to clot for 90 minutes in 1.5 mL microcentrifuge tubes. The clots were centrifuged to avoid access amount of unclot/liquid section. The enzyme activity was estimated by a change in the clot weight (Initial vs. final) by incubating it with an enzyme for 90 minutes. Standard was plotted with available streptokinase as STPase (Cadila Pharmaceuticals Ltd, Ahmedabad, India) with a 1500000 IU (International Unit). IU of streptokinase was measured as the ability to cause lysis of a fibrin clot via plasmin system *in vitro*. Standard was diluted as such that 100 µL of contains 3000 IU units. Total 8 Sets of 1.5ml micro centrifuged tubes containing clot blood were taken. Standard STPase dilutions from 1000-7000 IU were subjected to clot lysis in 7 micro centrifuged tubes and 1 tube with PBS for negative control. After 90 minutes, clot lysed blood was centrifuged to weigh the remaining clot. The initial clot was weigh was annotated as W1 and after treatment, it was annotated as W2. To calculate the percent (%) clot lysis following equation was used.

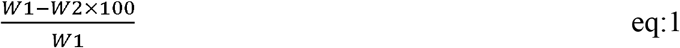

Standard was plotted for percent clot analysis vs IU. For two experimental samples, wild-type and mutant derived proteins, ammonium sulfate concentrated solution with an equal amount of protein 0.65 mg were subjected for clot lysis. Based on percent clot lysis their IU and specific activity were determined.

Fibrinolytic activity was estimated using a protocol described by [32]. A 100 µL of thrombin (10 NIH U/mL in 50 mM pH 7.2 sodium phosphate buffer) and fibrinogen from human plasma (5mg) was dissolved into 5 ml of 50 mM pH 7.2 sodium phosphate buffer. Thrombin and fibrin solutions were mixed into 2% agarose which was dissolved into 50 mM pH 7.2 sodium phosphate buffer and poured into Petri dish. Petri dish was allowed at 30 °C to form a fibrin clot layer. A 5mm well was created using a cup borer. The protein solution was added and allowed for clot lysis by incubating the plate at 37 °C for 12 hours. Plasmin (1000 IU) was used as a positive control. The zone of fibrin lysis was measured in mm.

### 2.10. Streptolysin heme release assay

*Streptococcus* species contains two types of streptolysins, streptolysin S which is oxygen liable, and streptolysin O is oxygen stable [33]. We used *S. equisimilis* contains the streptolysin S. Streptolysin activity was measured by percent release of heme during RBC lysis [34]. Substrate for Streptolysin S is human erythrocytes. A 10 ml blood was collected from a voluntary healthy individual (As per Institutional Animal Ethical Guideline). It was centrifuged at 3000 rpm for 30 mins at 4 °C. As shown in the supplementary file complete erythrocytes and plasma section was separated at 4 °C. Erythrocytes were washed in 10 PBS thrice to remove any impurities from the blood. Erythrocytes were dissolved in 10 ml PBS.1:20 diluted erythrocytes solution was used for studying Streptolysin activity using percent (%) heme release at A_540nm_. In a 1.5 mL micro centrifuge tube solution of erythrocytes was taken and ammonium sulphate was used to precipitate supernatant with equal amount of protein 0.65 mg. Tubes were incubated for different time interval ranging from 10 mins to 120 mins. On completion of each time interval tubes were centrifuged at 1000 rpm to settle down RBCs and only lysed RBCs heme section was present in supernatant that was determined by measuring OD at A_540nm_.

### 2.11. SagBCD complex computational modeling

X-ray crystal structure of SagBCD complex was not determined yet [7], however, it has shown structural similarity (**30%**) with *E. coli* McbBCD complex. Consequently, this complex was built by performing homology modeling using the McbBCD as template which is available in PDB (Protein data bank: 6GRI). We have used homology modeling methods to 3-D models of SagBCD complex (Schrodinger 2021.2 licensed suite).

## 3. Results

Streptolysin is hypothesized as a negative regulator by imparting instability of mRNA transcript of streptokinase. CRISPR-Cas9 based genome editing is required in *S. equisimilis* for enhanced levels of streptokinase expression by knockout/knockdown of Streptolysin S. Different gRNAs cassettes (four) were tried for the efficient knocking out. As discussed in the supplementary method section, golden gate assembly was used for gRNA insertion into pCRISPomyces-2 plasmid **(Supplementary figure 1**). Chimeric plasmids were maintained into *E. coli* top10F before inserting this plasmid into *S. equisimilis* to maintain efficient plasmid copy number for electroporation. The gRNA insertion was confirmed by Sanger sequencing **(Figure 1A-D)**. Detailed protocol from the preparation of chimeric plasmid to host insertion was added into supplementary section in figure 1. We have successfully constructed plasmid with desired gRNA.

**Figure 1:**
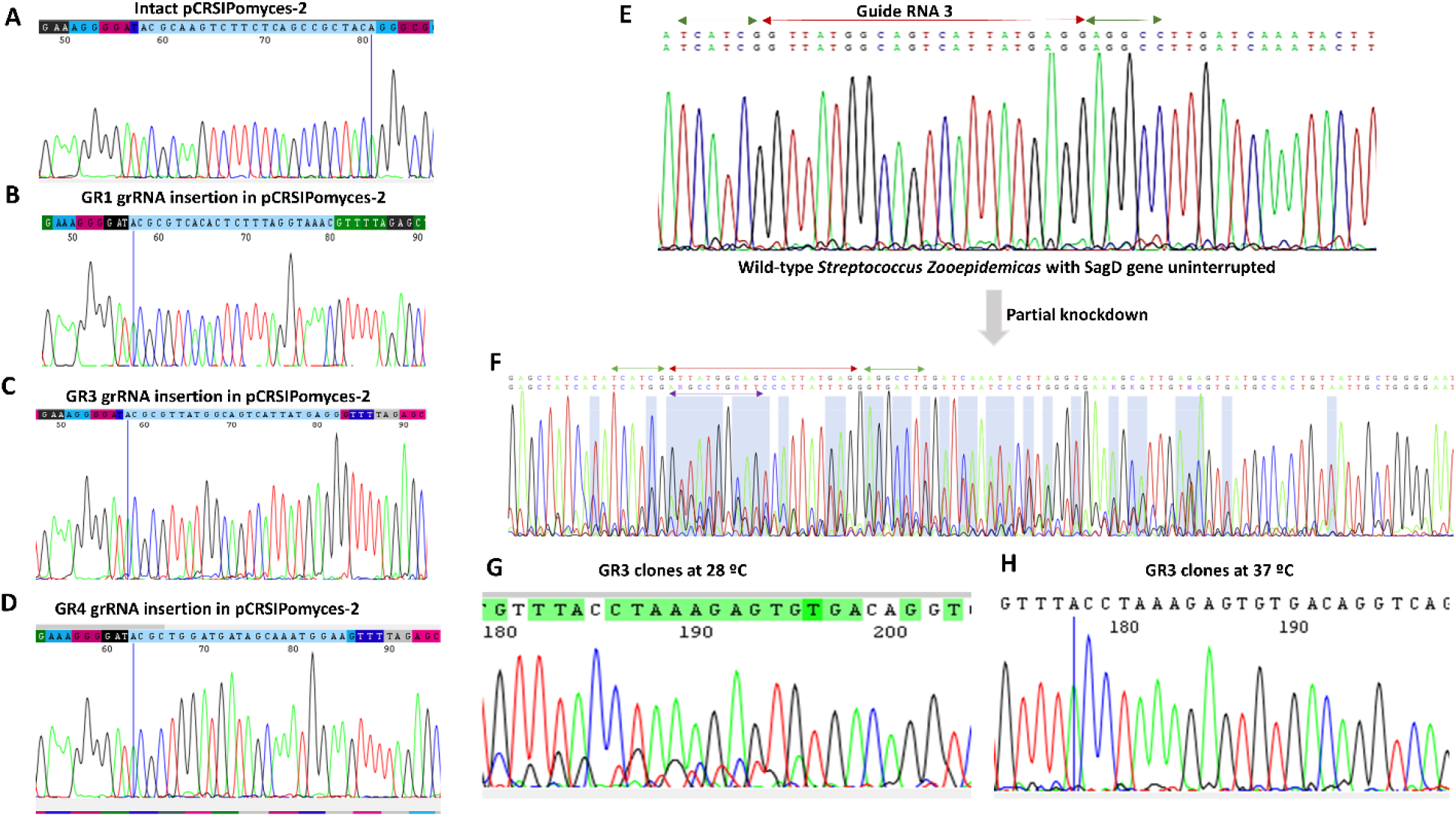
CRISPR-Cas9 system for knockout. **A: Intact** PCRISPomyces-2 plasmid. **B:** Incorporation of GR2 gRNA sequence into PCRISPomyces-2 plasmid. **C:** Incorporation of GR3 gRNA sequence into PCRISPomyces-2 plasmid. **D:** Incorporation of GR4 gRNA sequence into PCRISPomyces-2 plasmid. **E:** Native *sagD* gene sequence having intact/uninterrupted GR3 sequence. Red arrow showed the region covering GR3, green arrows were overlapped regions covering GR3. **F:** Poly-peak parser output for the clone sequences contained a mixed peaks for *sagD* gene. Purple arrow shows the sequence similar to the green arrow next to the GR3 sequence conforming mix population of clones having both edited and unedited *sagD* gene. **H:** Pcripomyces-2 was temperature-sensitive plasmids due to the present of PGF5. If clones having pCRISPomycespCRISPomyces-2 were grown at 37 °C they have curated the plasmid. If clones were grown at 28 °C they express the pCRISPmyces 2 plasmids and editing might occur. At 37 °C, no mix peaks were observed while at 28 °C, same sequence were there but mix peaks were observed. This may be possible because of the low expression of pCRISPomyces 2 plasmids for a complete knockout, which is leading to a mixing population having edited and unedited *sagD* gene.

### 3.1. Streptolysin knockout strategy using PCRISPomycesPCRISPomyces-2

The recombinant plasmid was prepared with four different gRNA cassettes. Figure 1A-D shows the insertion of gRNA sequences into pCRISPomyces-2 plasmid. pCRISPomyces GR1 and pCRISPomyces GR2 were first electroporated into *S. equisimilis* for Streptolysin knockout. PCripomyces was arranged as Cas9 expressed under control of rpsL promoter, which is 30S ribosomal protein S12 and gRNA is under gapdh (glyceraldehyde-3-phosphate dehydrogenase) promoter. Both promoters are proven strong promoters in wide varieties of microorganisms including bacteria, fungus algae and protozoa [20,35]. Screening of *S. equisimilis* clones having recombinant pCRISPomyces plasmid with *SagD* gene-specific gRNA constructs was subjected to grow at 28 °C in TSB media containing 50 µg/L Apramycine for 2-3 days for expression of these pCRISPomyces 2 system. Further these clones were screened using Sanger sequencing (genotypic) and altered hemolysis activity (phenotypic). However, we didn’t achieve any gene editing using these two gRNA cassettes GR1 & 2. Sequencing results were always followed one single pattern where region matching to gRNA has shown mixed peaks while outside region (other than 20bp guide RNA) have better coverage during DNA sequencing **(Figure 1F-H)**. This leads us to investigate sequencing results in further detail. We have further grown the same clone at two different temperatures 28 °C and 37 °C. As 28 °C temperature is a suitable temperature for pCRISPomyces-2 expression (37 °C leads to plasmid curation), continuous expression (at lower levels) of CRISPR array might be there in organisms which leads to the production of multiple strains with gene-edited and unedited region. Hence, mix peaks containing chromatograms was generated during sequencing **(Figure 1G-H)**. Poly peak parser helped to identify the second prominent sequence in the chromatogram from the region covering the gRNA specific to the *SagD* gene. Our analysis showed these second prominent sequences were lacking a 21 bp gRNA complementary region, which was shown as a purple arrow in **figure 1F**. Therefore, expected condition of constitutive expression of pCRISPomyces-2 plasmid with lower level is proved. These sequencing results allow us to conclude that there is a problem in the expression of higher transcripts of gRNA to bind specific genomic regions (*SagD*) for endonuclease activity **(Figure 1F)**. To overcome this issue, other gRNAs sequences were designed to have an efficient match up to 15-16 nucleotides out of 24 to *SagD* gene with a change in plasmid expression conditions have been shown in the material and method section. Those gRNAs sequences contained poly-A and poly T. The gRNA sequence binds to another region than the target region (off-targets other than *SagD* gene into *S. equisimilis* genome). Therefore, the binding of gRNA to a particular target has been divided and partial gene editing was observed. Graphical presentation for SagD gene editing is shown in **figure 2A-B**. Out of all 4 gRNAs, pCRISPomyces2-GR3 was successfully able to edit the genome in sagD region with 21 nucleotides deletion **(Figure 2C-D)**. Along with gRNA, another strategy for complete deletion of *SagD* gene, editing templet was created using overlap extension PCR **(Supplementary figure S3)**.

**Figure 2:**
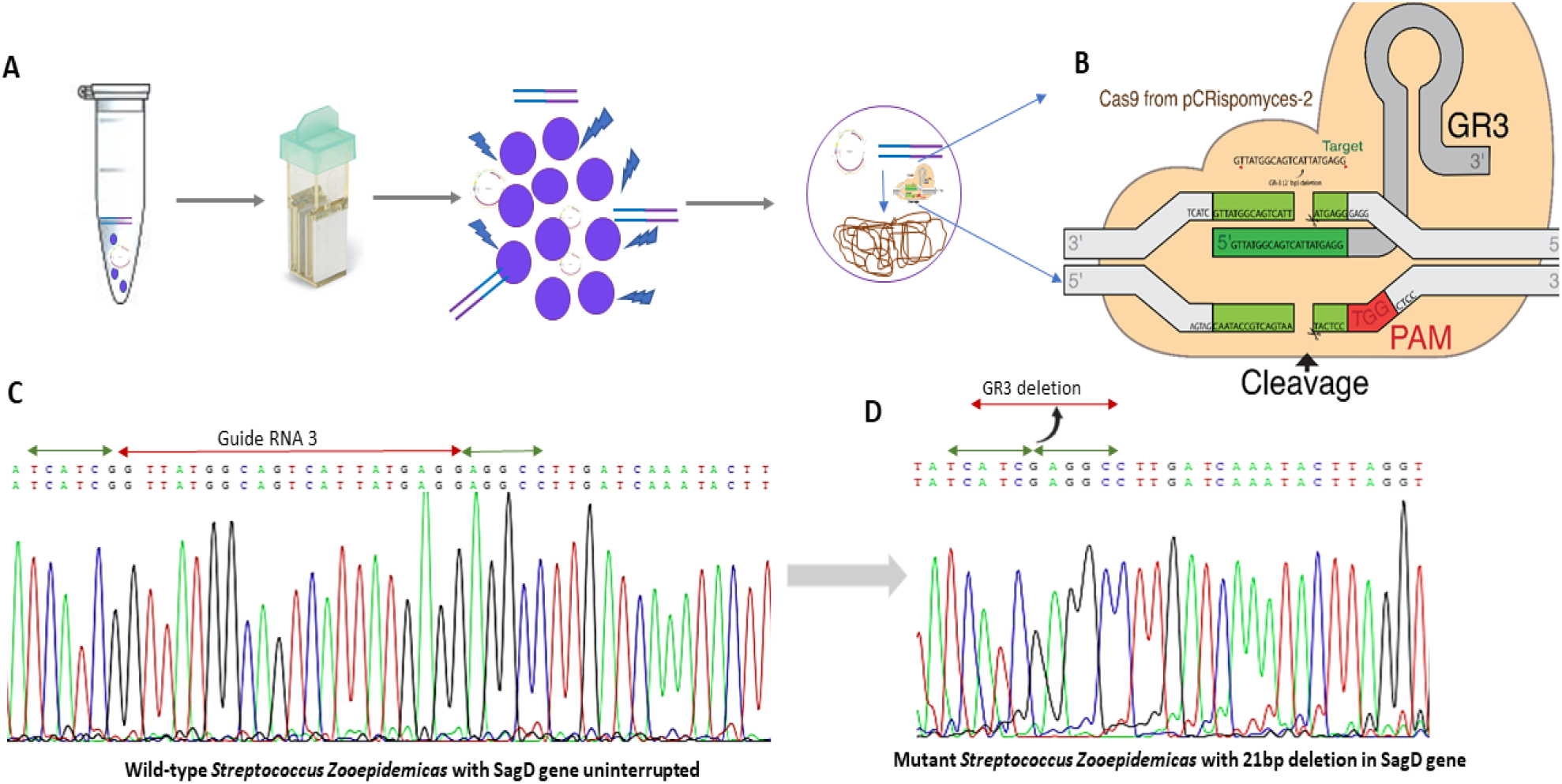
Gene editing of *SagD* gene using GR3-PCRISPomyces-2. **A:** Workflow showing an experimental design for inserting PCRISPomyces-GR3 with editing templet (SagC+E) using electroporation. **B:** Schematic diagram showing mechanism for gene editing using GR3. **C:** Native sagD gene without any gene editing. **D:** SagD gene sequence with missing 21 nucleotides complementary to GR3.

It was hypothesized that if gene editing template covering region *SagC* and *SagE* can be used then host cells can be used as a template to overcome endonuclease cut, hence during homology recombination, this *SagD* gene may be replaced by an editing template covering only *SagC* and *SagE* genes **(Supplementary figure S3B-D)**. However, complete deletion of *SagD* gene was not observed using the editing template. Clones having 21 bp deletion were used for further study to check the expression profile of both *SagD* and *streptokinase* genes. To investigate, whether 21 base pair deletion can disrupt the *SagA* post-translational modification or not. It was found that these 21 bp deletion/7 amino acid changes were altered the complex formation of the SagBCD complex, which might affect the *SagA* post-translational modification **(Figure 3A and Supplementary figure 4)**. SagBCD complex is prepared with E.coli MCB complex with Significant alpha and beta helices matches (30%) among both **(Supplementary figure 4A-B)**. Difference in root mean square deviation (RMSD) of wild-type and mutant SagBCD complex was 3.36 Å **(Figure 3A)**, which is significant.

**Figure 3:**
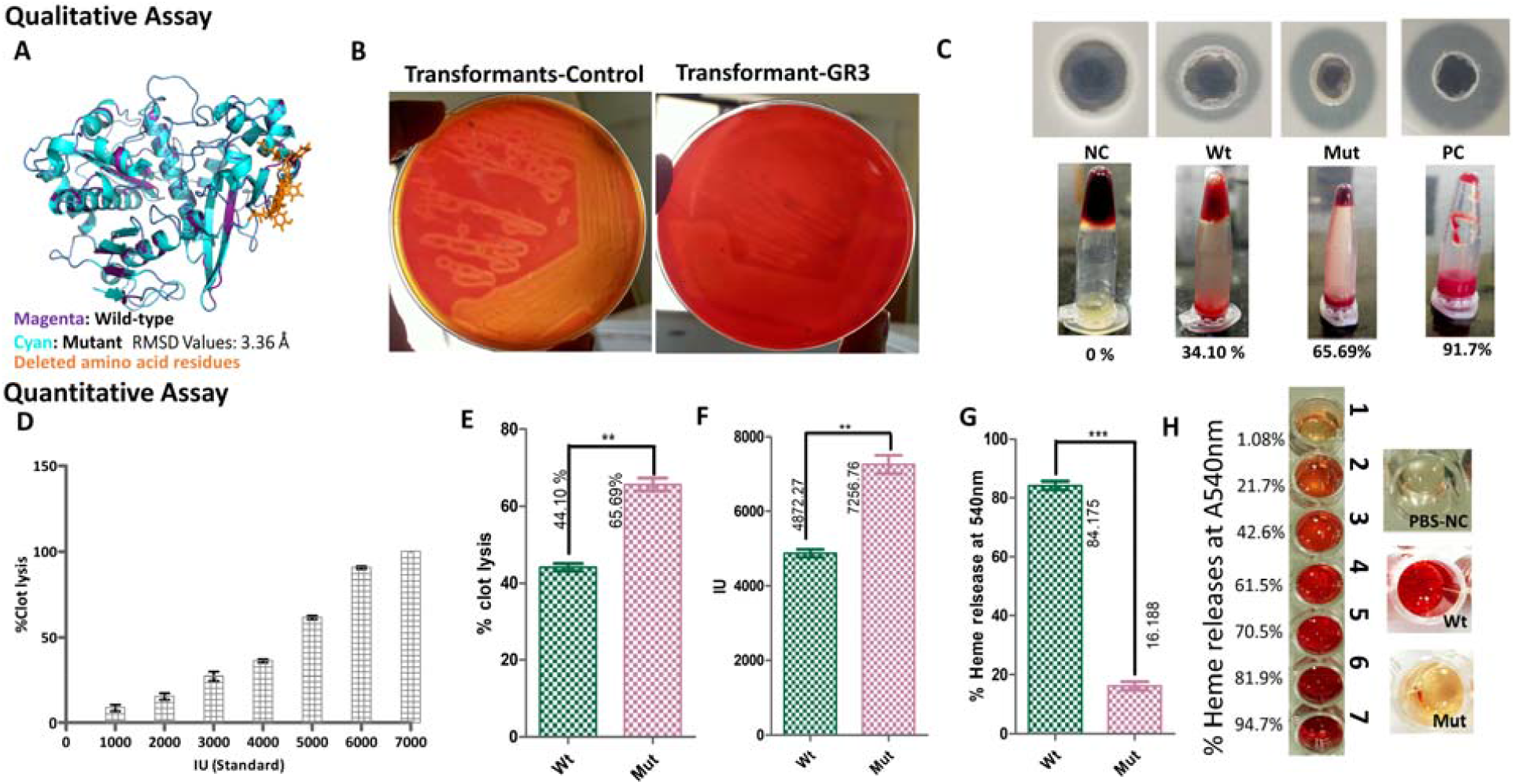
Quantitative and qualitative protein assays of streptokinase and Streptolysin. **A:** Structural superimposition wild-type and mutant SagBCD complex. The magenta color shows wild-type, cyan color shows mutant complex, and deleted amino acids are shown by orange color. **B**: wild-type and mutant strains were streaked on blood agar plates with equal 0.05 OD_600nm_, to show altered hemolytic activity. **C** Streptokinase clot lysis activity **C1:** Zone of fibrin lysis by streptokinase in the thrombin-fibrin agar plate. **C2:** Clot lysis assay for streptokinase. NC is abbreviated as a negative control, Wt (wild-type) and Mut (mutant) and PC is a positive control. **D:** Standard clot lysis assay of STPase for determination of IU units and specific activity of streptokinase. **E:** Percent clot lysis as compared in wild-type and mutant strain. **F**: Comparison of IU units within wild-type and mutant strain. **G:** Streptolysin activity represented using percentage heme release by lysis of erythrocytes. **H:** Change in percentage heme release measuring at A_540nm_ for Streptolysin.

**Figure 4:**
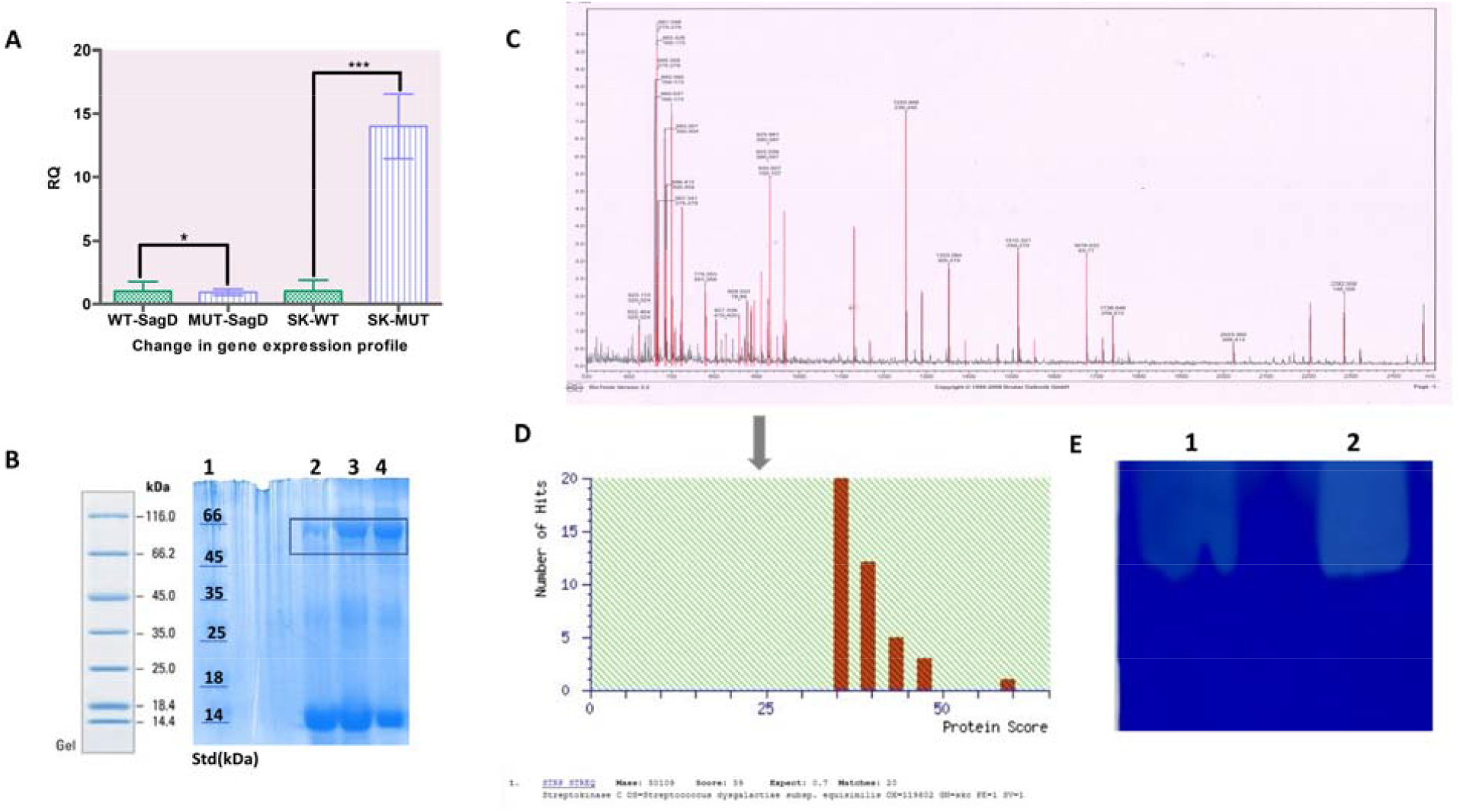
Determination of change in the expression profile of streptokinase. **A:** Real-time PCR data from gene expression panel showed expression of *SagD* and sks gene in wild-type and mutant. The green color bar showed gene expression data for wild-type and the purple color bar was shown data for mutant. **B**: SDS-PAGE analysis showed enhance expression of streptokinase at expected size ∼ 47kDa. lane 1: Protein ladder. lane 2: extracellular protein from wild-type. lane 3: Extracellular protein from mutant-GR3 strain. **C** and **D:** Peak pattern obtained using MALDI have matched with streptokinase protein. Red lines in peaks show 48% matching with streptokinase protein. MOSCOT analysis also showed streptokinase protein in 1^st^ hit. **E:** Fibrin Zymography showing high fibrin-thrombin clot lysis activity for mutant strain. Lane 1 shows the wild-type strain and lane 2 shows the mutant strain.

After validating genotypic change, phenotypic change was also corroborated using enzyme activity and altered hemolytic activity. As shown in **figure 3B**, GR3 edited clones have altered (reduced) hemolytic activity. In **Figure 3C**, wild-type gave zone, while mutant gave zone for clot lysis on a fibrin-agar plate. This qualitative change was further validated with quantitative clot lysis activity and hemolytic for streptokinase and Streptolysin, respectively. Blood is a common factor used in quantification for both enzymes, however, a substrate for each was different. It is important to separate individual substrates for both enzymes to remove any overlapping activity **(Supplementary figure 5S1)**. Blood was centrifuged at 4 °C for 30 min at 4000 rpm, Plasma and blood fraction were separated **(Supplementary figure 5S2)**. Plasma has a substrate for streptokinase which is plasmin and blood has a substrate for Streptolysin, which is erythrocytes (RBCs). *SagD* gene disruption leads to higher percent of clot lysis, leading to enhance expression of streptokinase. Percent clot lysis was increased from 44.10 ± 1.01 % to 66.69 ± 1.59 % after mutation (R^2^ = 0.985, P (0.0075) < 0.05) have been shown in **figure 3E**. Change in enzyme activity in IU was plotted with help of standard STPase **(Figure 3D)**. Wild-type strain showed 4872 ± 111.6 IU and mutant showed 7257 ± 175.6 IU (P < 0.05) in **Figure 3F**. After deletion streptokinase enzyme activity significantly increased due to the stability of mRNA transcript. Opposite to streptokinase, streptolysin activity seems to be significantly lowered, where the release of heme (hemoglobin) was not observed in GR3 edited clone **(Figure 3G-H)**. Percent heme release in wild-type strain is 84.18 ± 1.425 and mutant strain is 16.19 ± 1.068. Decreased the heme released in GR3 clone which was possible due to the altered streptolysin activity.

Enzyme activates of streptokinase and Streptolysin were shown to the enhance expression of streptokinase. However, expression may be due to the increased stability mRNA transcript specific to streptokinase (4). To validate enhance streptokinase expression, real-time PCR was carried out, where an increased number of *ska* mRNA transcripts may lead to higher fluoresces through sybergreen.

### 3.2. Change in the expression profile of streptokinase and Streptolysin

The change in the expression profile of both factors was validated using enzyme activity, which were further supported using SDS-PAGE for protein expression followed by RT-PCR. Strains were grown overnight and the next day, 0.05 OD 600nm was set up. Protein was harvested using ammonium sulfate and an equal amount of protein was loaded in well after carrying out protein estimation using Bradford assay. RNA isolation and cDNA synthesis as mention in section 2.7, equal concentration of cDNA was taken to perform gene expression assay. Change in gene expression was observed in SDS-PAGE were ∼ 50kDa band that was expected for streptokinase. To validate that these bands which indicate expression streptokinase enzymes, pmf (MALDI) and Zymography were carried out. According to [36], 21 bp deletion might not affect gene expression profile. RNA polymerase did not reduce the transcription of templates that have the deletion. Although, protein made up with deletion might have lost functionality. Streptolysin S protein was truncating mRNA transcript of Streptolysin using CoV-RS due which was sensitive to the ribonuclease and lowered the expression of streptokinase [2]. Altered Streptolysin was no longer able to suppress streptokinase expression due to which streptokinase mRNA was stable into a cell for a longer period. The study of fold change in gene expression data was analyzed using 2^ΔΔCT^ methods for relative quantification (RQ) (ABI PRISM® 7500 Sequence Detection System Software). Gene expression values were normalized with DNA gyrase endogenous control (control sample RQ = 1). *SagD* gene was shown a similar expression profile in both wild-type and mutant cases (t-test, P > 0.05). However, streptokinase expression was drastically increased in the mutant strain. To quantify the further change in expression of streptokinase in wild-type showed RQ = 1.03 while mutant showed RQ = 13.99 (t test, P value = 0.0009) **(Figure 4A)**. Fold change in streptokinase expression was 13.58 RQ. Gene expression data was completely shown into RT-PCR where enhance gene expression of streptokinase was observed.

Protein expression studies in wild-type and mutant were suggested to enhance the expression of streptokinase in the mutant **(Figure 4B)**. A higher protein band was observed in lane 3 which was streptokinase from mutant strain. It can be further validated with Western blotting but due to unavailability of antibodies for protein. It was not possible until availability of antibody. Therefore, we have MALDI-PMF and Zymography. To confirm these expression bands were subjected to MALDI-based pmf. MOSCOT analysis suggested streptokinase from *S. equisimilis* on the first hit, with query coverage of 48 % **(Figure 4D)**. As shown in **figure 4C** peaks matching to streptokinase were highlighted in red color. These data were completely supported the enhanced streptokinase expression that was also observed in RT-PCR. Zymography using fibrin and thrombin as a substrate was performed for streptokinase. Clear zones after Commasiee staining were obtained that suggested the fibrin clot degradation by streptokinase **(Figure 4E)**. In Figure 4E, lane 1 showed less clear zone of fibrin degradation as compared to lane 2 of mutant. Hence, enhance expression profile of streptokinase was validated using the standard protocols mentioned above. Here we successfully developed a CRISPR-Cas9 based gene editing model method for enhancing the expression of streptokinase.

## 4. Discussion

CRISPR-based techniques are extensively used in the fields from basic to applied biotechnology and medicine. We have used CRISPR-Cas9 genome editing in the prokaryotic organism for enhancing production of clinically important streptokinase. Streptokinase is an enzymes used as drug for treating myocardial infections, arteriovenous cannula occlusion, embolism, and deep vein thrombosis. It possesses a market value of USD40 million. Several methods were already available for increasing expression of streptokinase using wild-type *S. equisimilis* strain with various growth factors [37], plasmid based genetically engineered *E*.*coli* strain [38,39], and eukaryotic expression yeast [40,41]. Although maximum achieved production was 720mg/L streptokinase using *E*.*coli* BL21[9DE3] strain [39]. In the present study, we developed a model system in wild-type *S. equisimilis* strain using CRISPR-Cas9 mediated knocked out of streptolysin to enhance production of streptokinase. According to [2,42], streptolysin is a negative regulator of expression of streptokinase. Streptolysin is produced under the control of CoVRS/S system, which plays a key role for expression of virulence factors. *Streptococcus* possess small FasX RNA which imparts stability to streptokinase mRNA transcript and protects ribonucleases [2]. We targeted the *SagD* gene which forms a complex with SagBC to perform the post-translational modification to alter/knock down streptolysin S production.

*SagD* gene was mutated using pCRISPomyces-2 + gRNA. Protospacer specificity till 15-17 nucleotides is necessary for efficient binding to target without any off-target effect. BLASTn was performed while design of gRNA sequences to avoid any off-target effect. Sequences with minimum off-target effects were selected. A total 4 protospacer were used and GR3 was successfully able to edit *sagD* gene. Therefore, mutated *SagD* gene affects the streptolysin S production that was further validated using hemolysis assay. We observed a significant decrease in hemolytic activity **(Figure 3)**. Acceding to [43], the whole SagABCDEFGH cassette is important for the expression, post-translational modification, and transport of streptolysin. Deletion of any of sagABCDEFGHI genes can alter the SagA function as Streptolysin. Stable mRNA transcript of streptokinase was generated in *Streptococcus* after streptolysin alteration, mRNA pool-specific of streptokinase was also increased. An increase in streptokinase transcript was validated using SDS-PAGE followed by RT-PCR. The gene expression profile of streptokinase has significantly increased by 13.58 RQ (P-value < 0.05). This result supports the hypothesis showcasing stability of streptokinase mRNA transcripts leads to prolong expression of streptokinase in *S. equisimilis*. SDS-PAGE was shown (in **figure 4B**) clearly indicates the enhanced expression of streptokinase (50kDa). Although it was concentrated crude supernatant, not purified protein. Here native strains were used for expressed protein was in a native form without having any affinity Tag like histidine. To validate the SDS-PAGE expression profile, and MALDI. Protein band of ∼50 kDa were excised and subjected to pmf-MALDI, with results in more than 40% hit specific to streptokinase in MOSCOT search. Hence SDS-PAGE figure print of increased streptokinase expression was conformed. To further validate, an equal concentration of streptokinase was used for substrate-specific Zymography.

Studies on the expression of streptokinase with different expression vectors/strategies and the host are available. Some have used *Streptococcus agalactiae* with nonspecific ethyl methanesulfonate (EMS) based chemical mutagenesis [44], *Streptococcus sanguis* produced lowered molecular mass containing streptokinase [45], *B. sublitis* with C-terminally processed streptokinase[46]. Relative comparison of streptokinase activity in different hosts was as follows, *Saccharomyces pombe* (2450 IU), *Streptococcus equisimilis* (100–150 IU), *Pichia pastoris* (3200 IU) and *E. coli* (1000–1500 IU) [44,47–49]. All these studies observed that streptokinase degradation of matured streptokinase 47kDa to inactivated 44kDa [45,46,50]. In this present study, we developed model system for streptokinase expression without requirements of inducer, antibiotic, and optimized media in native strain. Our engineered strain can be produced streptokinase in a cheaper cost. In native strain, streptokinase degradation was not observed which is another advantage. As shown in Figure 4E, lane 1 was wild-type with less zone of fibrin clot lysis and lane 2 contains a wild-type streptokinase enhanced zone of fibrin clot lysis. Clot-buster thrombolytic streptokinase activity was measured in terms of IU sing 1.5 million IU standard STPase. Fold increase in streptokinase-specific activity in terms of IU per ml was 1.12. Fold change in enzyme activity of streptokinase from wild-type and mutant was 1.48 significant with a P value less than 0.05.

CRISPR-Cas9 system helps us to alter *sagD* gene which is an alternative approach for enhanced expression of streptokinase. Here native strain can be used for industrial streptokinase production where its virulence is altered and desired product expression is increased. No inducer is required as the product is generated by native strains only. CRISPR-based gene editing can be useful for the production of efficient microorganism models for production of industrially important products.

## 5. Conclusions

Streptokinase is an industrially important enzyme and used as a drug for breaking down of blood clotting in some cases of myocardial infarction (Heart attack), pulmonary embolism, and arterial thromboembolism. In this study, we have successfully designed CRISPR-Cas9 based system for *SagD* gene (inhibitor for streptokinase expression) knock out in wild-type of *S. equisimilis* strain. We have found 13.58-fold relative quantification of streptokinase in knocked out strain as compared to wild-type strain. Our CRISPR-Cas9 based approach can be used in other microorganisms for genome editing for other industrially important therapeutic production.

## 6. Highlights

- Streptokinase is an important enzyme used as a drug for breaking down of blood clotting.
- *S. equisimilis* strain was used for expression of streptokinase
- CRISPR-Cas9 system was developed for genome editing of sagD gene
- We observed 13.58-fold higher streptokinase in knocked out strain as compared to wild-type strain, also fold change in enzyme activity is 1.48 (t-test, P > 0.05)

## 7. Acknowledgments

Authors acknowledge Dr. Poonam Bhargava and Dr. Manju shri Chaudhary for their support in selecting plasmid compatible for this study. Authors acknowledge Gujarat State Biotechnology Mission, DST, and Government of Gujarat, India for financial support to carry out this work.

## 8. Author’s contribution

AC, SV, MJ, CGJ conceptualized the study. AC designed the experiments. AC performed the experiments. AC, MJ, AL, VS, and CGJ wrote, edited, revised and supervised. All authors reviewed, edited and approved manuscript.

## 9. Conflict of interest

Authors declared no conflict of interest.

---

**Table 1:**
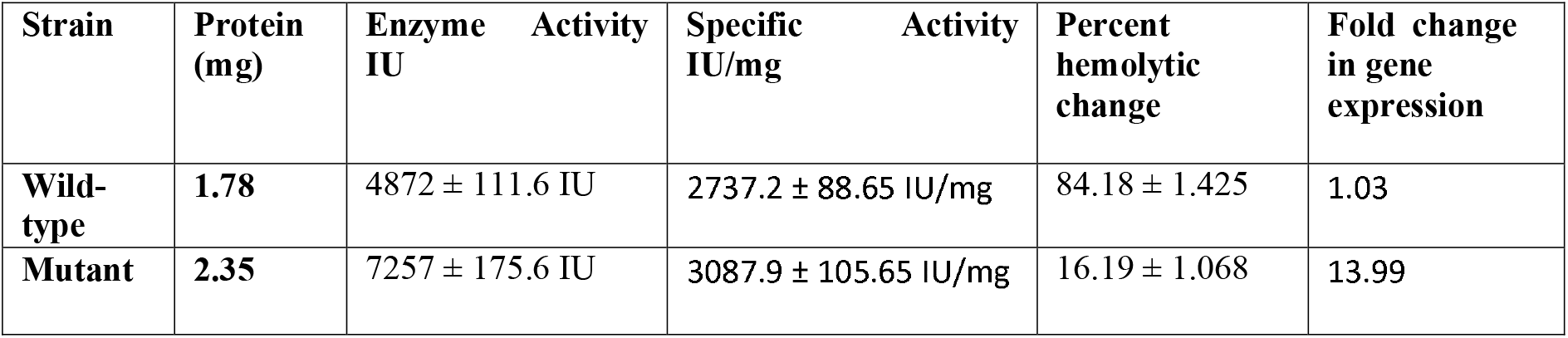
Summary of streptokinase expression in wild-type and mutant strain of *S. equisimilis*.

## Notes

### Competing Interest Statement

The authors have declared no competing interest.

