## Supplemental figures for "CRISPR-Cas9 mediated knockout of SagD gene for overexpression of streptokinase in *Streptococcus equisimilis*": Supplementry CRISPR.docx

**Supplementary section:**

**1. pCRISPomyces-2 expression and screening of clones for GR3 inserts.**

Golden gate assembly protocol was followed for insertion of four guide RNA sequences to the pCRISPomyces-2 plasmid. Supplementary figure 1, explains the elaborated information about chimeric plasmid preparation.

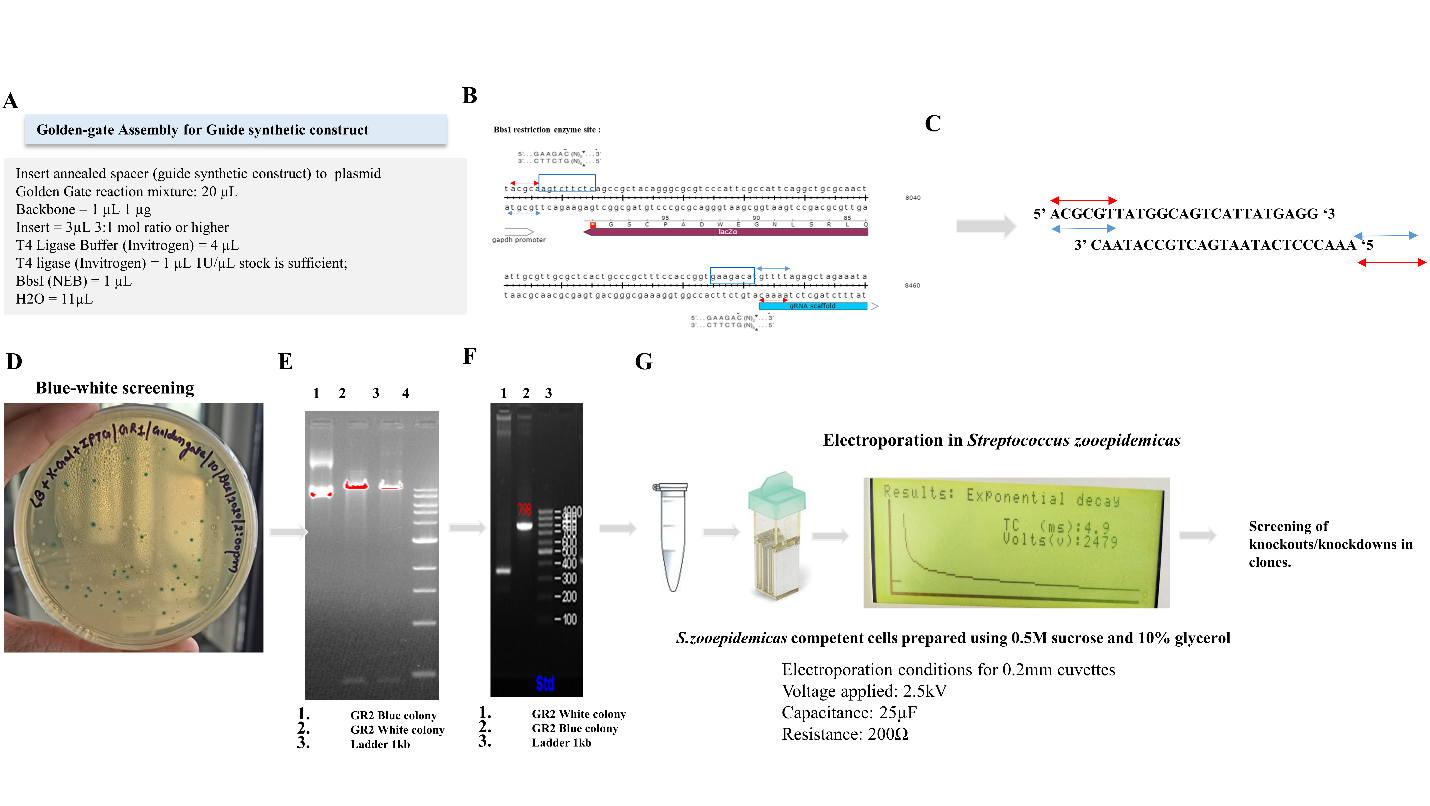

**Supplementary figure S1:** Golden gate assembly protocol for chimeric plasmid preparations. **A:** Golden gate reaction mixture where Bbs1 and T4-DNA ligase were added into a single tube and PCR conditions favoring simultaneous cut and ligation were applied**. B:** Two sites of Bbs1 were present in pCRISPomyces-2 plasmid to produce staggered overhangs. **C:** Guide RNA sequence**s** with ACGC and CAAA overhang were designed. These overhangs were complementary to Bbs1 restriction sites for better ligation. **D:** After heat-shock transformation of the chimeric pCRISPomyces-2 plasmid into *E.coli* top 10F’ cells, blue-white screening was used to select the plasmid containing guide RNA construct. **E:** White transformation was further validated using Bbs1 restriction where the blue colony will yield a single 10 kb product and Transformants (white colony) will yield intact supercoiled, nicked plasmid as they didn’t possess bbs1 restriction site anymore. **F:** clones were also validated using pCRISPmyces-2 F & R primers, where blue colony having intact LacZ gene will yield ~800bp band and the white colony will yield only ~300 bp due to deletion of lacZ gene showing guide NRA insertion. **G:** *Streptococcus equisimilis* was washed with sucrose and 10 % (V/V) glycerol and further transformed using 0.2 mm electroporation cuvettes of Biorad gene pulser.

**2. PCRISPOMYCES 2 expression in streptococcus equimilus**

*Streptococcus equisimilus* is gram-positive organism, which is difficult to get electroporated using chemical transformation, hence electroporation was used. Clones were conformed using pCRISPomyces specific pair of primers pCRISPomyces-2_GR.

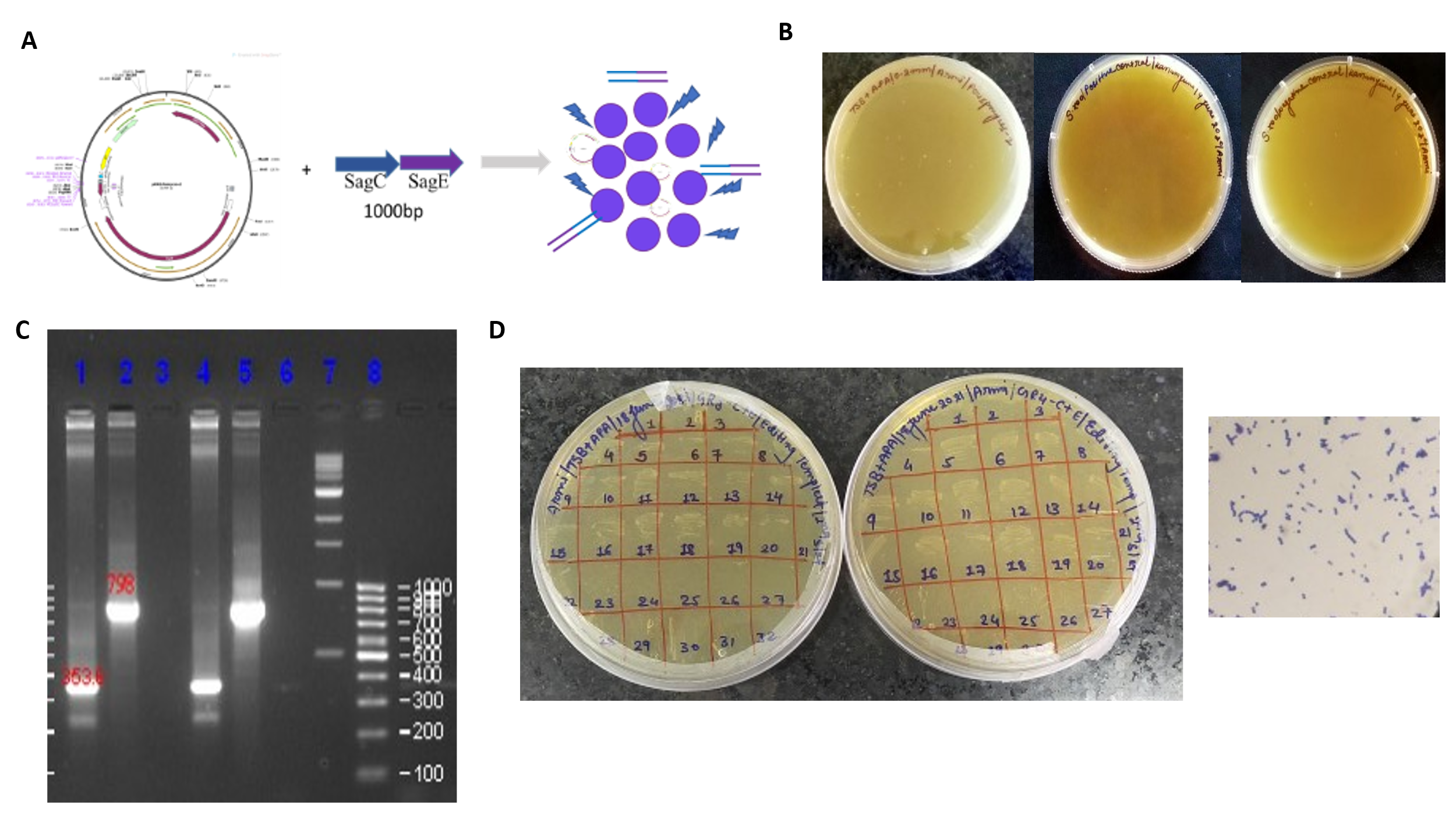

**Supplementary Figure 2:** PCRISPmyces-2 insertion in host **S2A:** Schematic representation of pCRISPmyces-2 plasmid and strategy for electroporation Streptococcus equimilus cells. **S2B:** transformation of pCRISPmyces-2 into streptococcus equimilus using electroporation. **B1:** GR3 transformants **B2:** Positive control **B3:** Negative control. **S2C:** PCR amplification of for conformation of pCRISPmyces-22 plasmid using pCRISPomyces2_GR. Control cells have intact lacZ gene so amplification product is 800bp and GR3 clones with deletion of LacZ gene it will show around 350 bp amplification. **S2D:** Random colonies were picked up and transfer in to TSB + Apramycin containing medium. Gram staining was always performed to conform purity of culture.

**3. Gene editing template for *SagD* gene knockout**

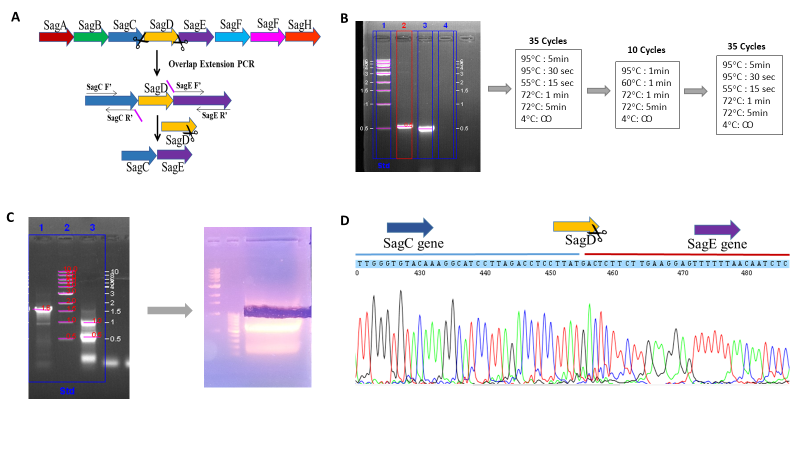

**Supplementary figure S3:** Editing templet strategy for complete removal of SagD gene. **S3A:** cassette for streptolysins S biosynthesis is shown. Gene A is forming pre-streptolysin which requires post translational modification using SagBCD complex. SagEFG transport function for streptolysin. Overlapping primers of sagC and sagE were designed as mentioned in material and method section X. SagC’ R and Sag E’ F have complementary region which will be used for overlap extension pcr. **S3B**: PCR amplification using OEPSagC and OEPSagE, was performed. Lane 1: shows 1kb NEB ladder. Lane 2: 500bp product of OEPSagC. Lane 3: 500bp product if OEPSagE. Lane 4: NTC (non templet control). First pcr for OEPSagC and OEPSagE was performed, second pcr with complementary annealing and extension condition for OEPSagCR and OEPSagEF was performed. Product of pcr 2 was further used to performed pcr 3 amplification from OEPSagF and OEPSagER. S4C: Product of pcr 3 is editing templet. Lane 1: sagD gene which is 1.67 Kb, Lane 2: 1kb NEB ladder. 3: 1kb is editing template generated using OEPSagCF and OEPSagER primers. To overcome issue of nonspecific amplification agarose gel extract was done for 1 kb editing templet. **S3D**: Sequencing was performed for sagC and SagE gene containing editing where missing portion was sagD gene was conformed. Quality of chromatogram was compromised due to overlap extension PCR, but editing templet conformation can be done.

**4. Computational Analysis for SagBCD complex**

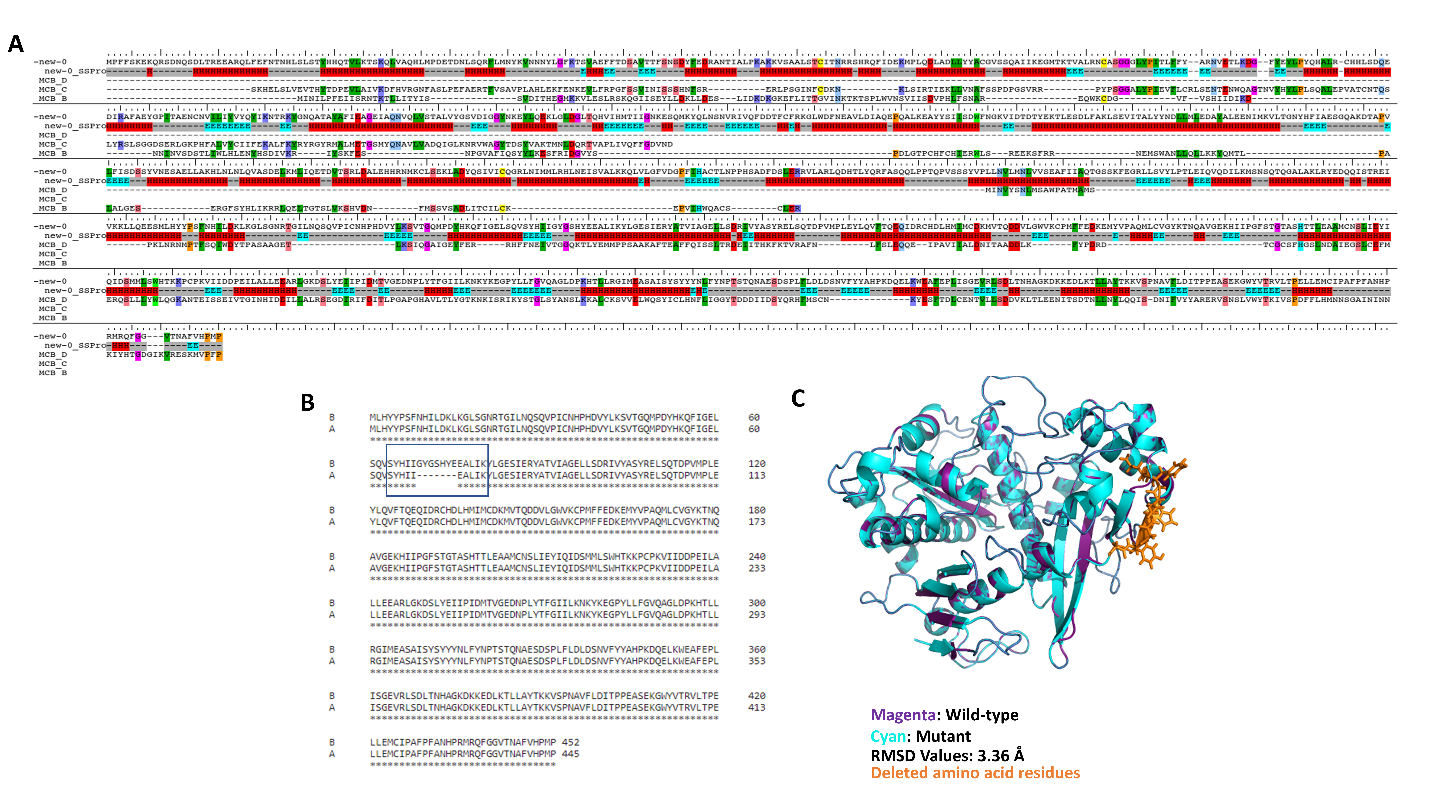

**Supplementary figure 4: S4A:** sequence alignment of SagBCD complex with wild-type e.coli MCB complex is shown here. Red colored region is alpha helix and blur colored region is beta sheets. **S4B:** Alignment of wild-type and mutant sequence from sagD gene for conformation of 7 amino acid deletion. **S4C:** In figure S4C structural alignment of homology modelled structure of wild0type and mutant sagD gene is shown.

**5. Enzyme activity**

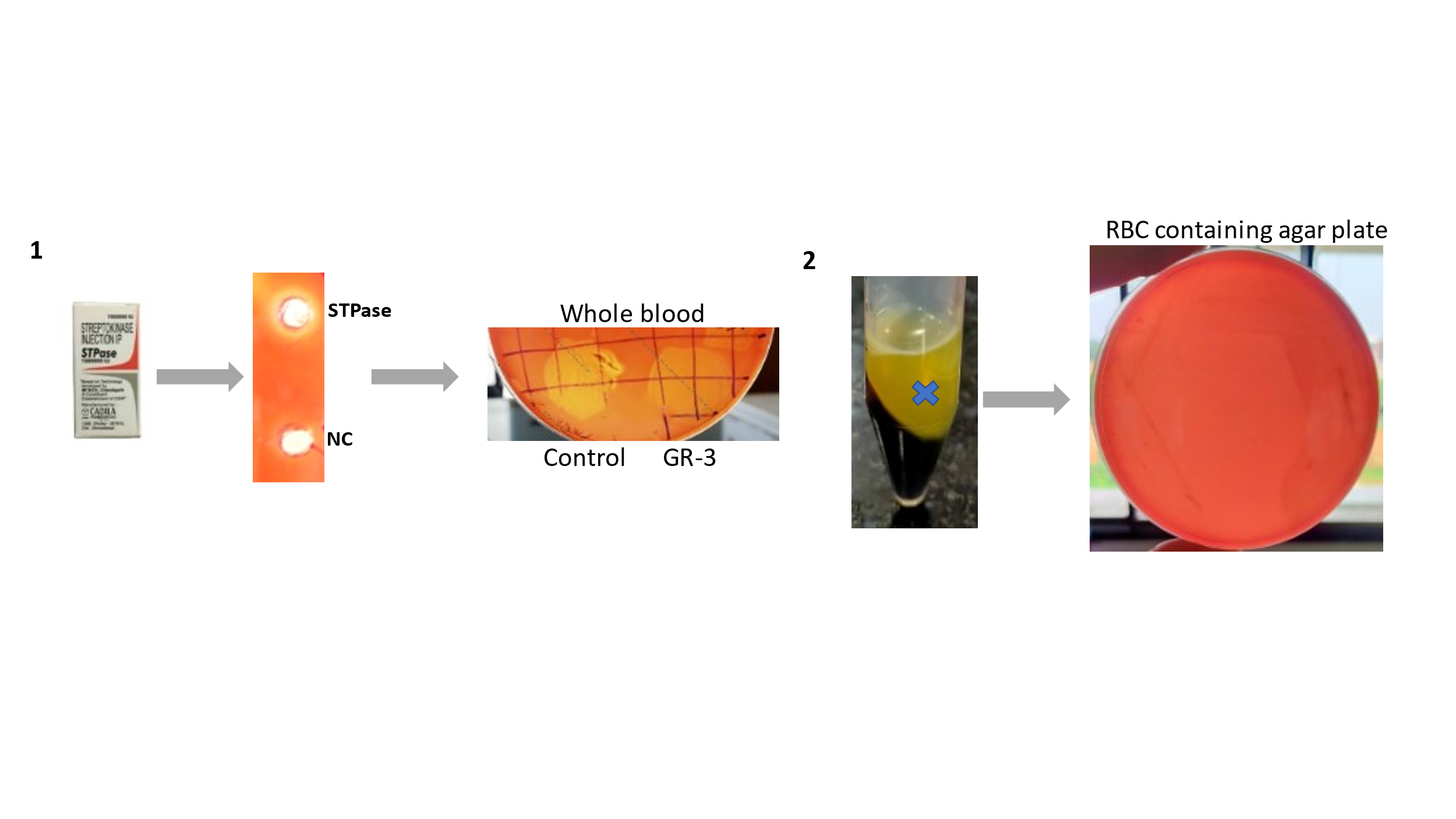

**Supplementary Figure S5: 5S1:** Streptokinase standard in available as STPase, produced by cadila pharmaceuticals. STPase gave partial zone on blood agar plate, reason behind it was presence of euglobin fraction in blood. Euglobin is substrate for streptokinase. Hence partial zone was observed. Streptokinase activity is measured by plasma faction and streptolysine activity is measured by erythrocytes. To separates substrate for both factors blood was subjected to centrifuge at 4000 rbc for 30 mins to separate out both sections. **5S2:** Erythrocytes were dissolved in pbs and used in rbc agar plate formation. GR-3 clones were showing no zone of hemolysis in rbc agar plate, which supports the altered streptolysine activity.

**Supplementary Table 1: List of all the organisms used in the study.**

| **Sr no.** | **Name of the organism** | **Selection** |
| --- | --- | --- |
| **1.** | *E. coli Top10F’* | NA |
| **2.** | *E. coli Top10F’-*pCRISPomyces2 | Apramycin |
| **3.** | *E. coli Top10F’-*pCRISPomyces2-GR1 | Apramycin |
| **4.** | *E. coli Top10F’-*pCRISPomyces2-GR2 | Apramycin |
| **5.** | *E. coli Top10F’-*pCRISPomyces2-GR3 | Apramycin |
| **6.** | *E. coli Top10F’-*pCRISPomyces2-GR4 | Apramycin |
| **7.** | *S. equimilus* | NA |
| **8.** | *S. equimilus* *-*pCRISPomyces2 | Apramycin |
| **9.** | *S. equimilus* *-*pCRISPomyces2-GR1 | Apramycin |
| **10.** | *S. equimilus* *-* pCRISPomyces2-GR2 | Apramycin |
| **11.** | *Streptococcus equimilus* *-* pCRISPomyces2-GR3 | Apramycin |
| **12.** | *Streptococcus equimilus* *-* pCRISPomyces2-GR4 | Apramycin |

**Supplementary Table: 2 List of primer and gRNAs sequences used in this study.**

| **Sr no.** | **Name of the Primer Pairs** | **Forward sequence(5’-3’)** | **Reverse Sequence(5’-3’)** |
| --- | --- | --- | --- |
| 1. | SagD gene | ACAAGGGGCTCTGGCAAAAT | AAGAGCACAGCCAAGCTCAA |
| 2. | Streptokinase gene | ATGTTGACTTCAAAAAGAC | TTATTTTTGCAGGTCTT |
| 3. | pCRISPomyces2_GR | ACGGCTGCCAGATAAGGCTT | TTCGCCACCTCTGACTTGAG |
| 4. | SKC_RT | ATACATCTTGACGGGTCAGG | AAGAGACCCTGCTGCCAT |
| 5. | SagD_RT | GACAGCCTCTCATACAACAC | AGCGGATTATCCTCTCCAAC |
| 6. | GR1_SagD | ACGCGTCACACTCTTTAGGTAAAC | AAACGTTTACCTAAAGAGTGTGAC |
| 7. | GR2_SagD | ACGCGGACTTAGTGGTAACCGAAC | AAACGTTCGGTTACCACTAAGTCC |
| 8. | GR3_SagD | ACGCGTTATGGCAGTCATTATGAGG | AAACCCTCATAATGACTGCCATAAC |
| 9. | GR4_SagD | ACGCTGGATGATTAGCAAATGGAA | AAACTTCCATTTGCTAATCATCCA |
